# Manipulation of Photosensory and Circadian Signalling Restricts Developmental Plasticity in Arabidopsis

**DOI:** 10.1101/2024.06.17.598603

**Authors:** Martin William Battle, Scott Fraser Ewing, Cathryn Dickson, Joseph Obaje, Kristen N. Edgeworth, Rebecca Bindbeutel, Rea Antoniou Kourounioti, Dmitri A. Nusinow, Matthew Alan Jones

## Abstract

Plants exploit developmental plasticity to adapt their growth and development to prevailing environmental cues. This developmental plasticity provides a selective and competitive advantage in nature but is obstructive during large-scale, intensive agricultural practices since economically important traits (including vegetative growth and flowering time) can widely vary depending on local environmental conditions. This prevents accurate prediction of harvesting times and produces a variable crop. We sought to restrict developmental plasticity by manipulating signalling systems that govern plants’ responses to environmental signals. Mathematical modelling of plant growth and development predicted a reduction in plant responses to changing environments when circadian and light signaling pathways were manipulated. We tested this hypothesis by utilising a constitutively-active allele of the plant photoreceptor phytochromeB, along with disruption of the circadian system via mutation of *EARLY FLOWERING3.* We found that the combination of these manipulations produced plants that are less responsive to light and temperature cues. These engineered plants have uniform vegetative growth and flowering time and demonstrate how developmental plasticity can be limited whilst maintaining plant productivity. This has significant implications for future agriculture in both open fields and controlled environments.

## Introduction

Developmental plasticity enables plants to adapt to micro-niches within their environment but is problematic in modern agriculture which benefits from uniform and predictable growth and reliable time to harvest. In addition to experiencing daily and seasonal climatic differences, plants respond to light and temperature signals differentially dependent upon time of day (Millar, 2016). Photo- and thermo-sensors work in combination with the circadian system which provides an internal timing reference relative to dawn and dusk (Sanchez et al., 2020, Kerbler and Wigge, 2023). The plant circadian system continually integrates light and temperature as entrainment signals to modulate development (Webb et al., 2019). A suite of photoreceptors including phytochromes (phyA through phyE), cryptochromes (cry1-3), zeitlupe (ZTL), and UVR8 each integrate light signals into the circadian clock (Somers et al., 2004, Fehér et al., 2011, Somers et al., 1998, Sanchez et al., 2020). Photoreceptors integrate with the circadian system at multiple levels, with phytochromes physically interacting with EARLY FLOWERING3 (ELF3), a protein which is essential to maintain circadian rhythms (Covington et al., 2001, McWatters et al., 2000, Thines and Harmon, 2010, Huang et al., 2016a). Both phyB and ELF3 have been shown to be responsive to temperature in addition to their roles in photoperception and the circadian system and are therefore crucial components of the sensory system that enable plants to adapt to prevailing environmental conditions (Jung et al., 2020, Jung et al., 2016, Legris et al., 2016).

## Results and Discussion

### Modelling refines our understanding of light input into the circadian system

Decades of research suggest that phyB- and ELF3-signalling pathways are genetically separable, although multiple lines of evidence demonstrate a functional interaction between these signalling pathways (Reed et al., 2000, Kolmos et al., 2011, Covington et al., 2001, Jung et al., 2016, Legris et al., 2016, Liu et al., 2001, Yu et al., 2008, Nieto et al., 2022). Mathematical modelling of these interactions highlights the central contributions of phyB and ELF3 towards key aspects of development such as seedling establishment and flowering time (Chew et al., 2022, Seaton et al., 2015). Disruption of ELF3 function induces consistently early flowering, yet imposes an etiolated phenotype that is reproduced by the Arabidopsis Framework Model [FMv2; Fig. S1A-C, (Chew et al., 2022, Seaton et al., 2015)]. Since increased phyB activity (either through over-expression or inclusion of a constitutively-active Y276→H [YHB] allele) promotes photomorphogenesis via post-translational regulation of PIFs we expected that YHB would be epistatic to *elf3* with regards photomorphogenesis (Hajdu et al., 2015, Su and Lagarias, 2007, Wagner et al., 1991). FMv2 aligned with our hypothesis that increased phyB signalling in the absence of ELF3 would limit hypocotyl growth whilst retaining an early flowering phenotype (Fig. S1A-C).

Plants expressing *YHB* maintain robust circadian rhythms in constant darkness compared to wild type, although it remains unclear how phyB-initiated signals are integrated into the circadian system (Jones et al., 2015, Huang et al., 2019). We examined two alternate hypotheses to apply constitutive phyB signalling into the circadian module of FMv2 (Fig. S1D). Initially we examined whether constitutive phyB signalling acted by promoting light-induced gene expression within the model, as well as repressing COP1 accumulation (Fig. S1D). This ‘global phyB effect’ was unable to reconstitute YHB-mediated circadian rhythms in FMv2 after transfer to constant darkness (Fig. S1E). Interestingly, work examining dawn-induced gene expression suggests that photoreceptor activation is insufficient to promote transcript accumulation (Balcerowicz et al., 2021). Removal of light-activated gene expression from our YHB simulation provided a ‘COP1 only’ variant that retained circadian rhythms in constant darkness but showed similar behaviour to wild type and therefore remained inconsistent with previous experimental data [Fig. S1D, S1F; (Pokhilko et al., 2012, Fogelmark and Troein, 2014, Jones et al., 2015, Huang et al., 2019)].

This inconsistency encouraged us to examine alternate circadian models. The F2014 circadian model revises the FMv2 circadian module to include refined waves of transcriptional repression based on experimental data (Fogelmark and Troein, 2014). The resultant ‘FMv2+F2014’ model recapitulated YHB-mediated retention of circadian amplitude compared to damping in wild type (Fig. 1). The effect of YHB was apparent in both ‘global’ and ‘COP1 only’ approximations of YHB, although again the ‘COP1 only’ variant matched the experimental luciferase data more closely [Figs. 1B-D; (Huang et al., 2019, Jones et al., 2015)]. We next examined how disruption of *ELF3* was predicted to affect constitutive phyB signalling. Both FMv2 and FMv2+2014 models predict *elf3* will be epistatic to *YHB* with regards circadian rhythmicity [Fig. 1 and S1; (Thines and Harmon, 2010, McWatters et al., 2000, Covington et al., 2001, Huang et al., 2016a)].

**Figure 1.**
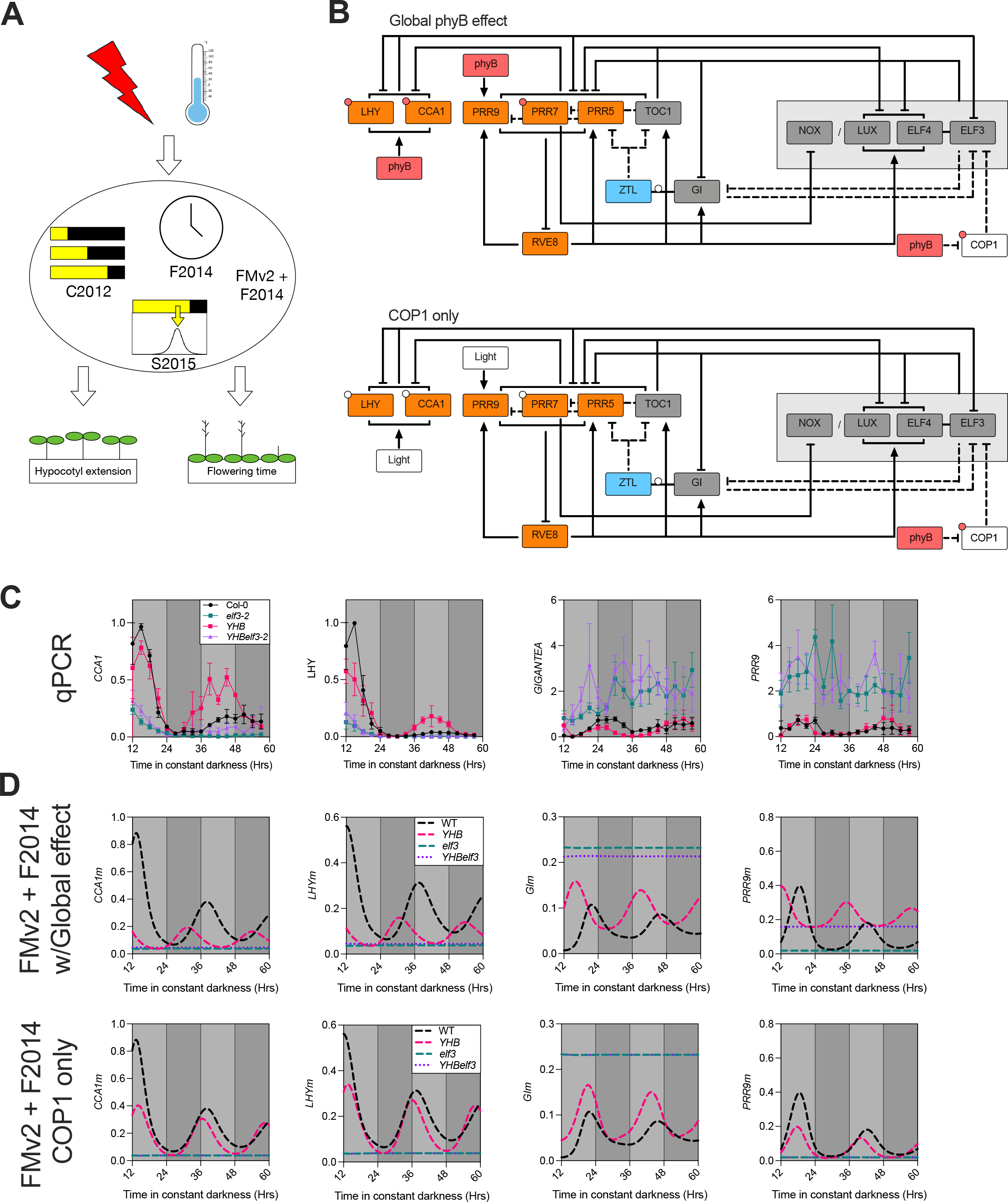
Manipulation of phyB and ELF3 activity is expected to reduce developmental plasticity in plants (A) Cartoon of the revised Arabidopsis Framework Model v2 including F2014 circadian model (FMv2+F2014) (Chew et al., 2022) **(B)** PhyB signalling into the circadian system was modelled via two hypotheses. The ‘Global phyB effect’ variant (upper) proposes that activated phyB is sufficient to induce light-activated gene expression in the circadian system in addition to enabling degradation of COP1. The ‘COP1 only’ variant (lower) restricts the effect of phyB activation solely to the turnover of COP1. In both cases, stability of ZTL and GI is regulated independently since this is a blue light- mediated effect effect (Kim et al., 2013). Circadian model adapted from F2014 (Fogelmark and Troein, 2014). Post-translational regulation by light is shown by small white circles. Small red circles indicate post-translational regulation induced by phyB. **(C)** Accumulation of *CCA1*, *LHY*, *GIGANTEA*, and *PRR9* in constant darkness. Plants were entrained in 12:12 light:dark cycles for 12 days before being transferred to constant darkness at dusk (ZT12). Tissue was sampled every 3 hours at the timepoints indicated. Data presented is the average of three independent biological replicates, and is presented relative to accumulation of *APA1, APX3,* and *IPP2* transcripts. Error bars indicate SEM. **(D)** Modelled accumulation of *CCA1m* (*CCA1* mRNA)*, LHYm* (*LHY* mRNA), *GIm* (*GIGANTEA* mRNA), and *PRR9m* (*PRR9* mRNA) in constant darkness. Light grey bars demonstrate subjective day in constant darkness.

### The combination of *YHB* and *elf3* alleles restricts daily patterns of gene expression

We next sought to reproduce these predictions *in planta* by introducing the *YHB* allele into *elf3* (Hu et al., 2009, Su and Lagarias, 2007). This allowed us to assess whether *YHBelf3* seedlings had phenotypes aligned with our modelled predictions, with the ultimate goal of minimising developmental plasticity in plants (Fig. 2). *In vivo,* neither constitutive expression of *YHB* [*35S::YHB* (*elf3-1 phyb-9*)] nor expression of *YHB* driven by the endogenous *PHYB* promoter [*PHYB::YHB* (*elf3-2); YHBelf3-2*] were able to maintain circadian rhythms of *CCA1*-driven bioluminescence in constant darkness, with only 15% of *YHBelf3-2* lines being assessed as rhythmic [Fig. 2A-B and Fig. S2A-B; (Jones et al., 2015, Huang et al., 2019)]. qRT-PCR analysis of candidate genes (including *CCA1, LHY, GIGANTEA,* and *PSEUDORESPONSE REGULATOR9; PRR9*) confirmed the loss of circadian rhythmicity in *YHBelf3-2* lines compared to *YHB* (Fig. 1C). Interestingly, mis-regulation of these candidate circadian transcripts fell into two groups; *CCA1*/*LHY* [whose promoters are solely bound by phyB; Fig. S3; (Ezer et al., 2017, Jung et al., 2016)] and *GI/PRR9* [bound by both phyB and ELF3; Fig. S3; (Ezer et al., 2017, Jung et al., 2016)]. For each transcript, accumulation patterns over time were consistent in *elf3-2* and *YHBelf3-2* seedlings (Fig. 1C). Although the FMv2+F2014 model aligned with experimental transcript accumulation for *CCA1, LHY*, and *GIGANTEA*, we were interested to note that the FMv2+F2014 model predicted *PRR9* mRNA to damp to basal levels in *elf3* and *YHBelf3* plants (Fig. 1D). This contrasts our experimental data which demonstrates elevated (and arhythmic) *PRR9* accumulation in *elf3*-2 and *YHBelf3-2* plants (Fig. 1D). Such data demonstrate that ELF3 is necessary to retain circadian rhythms yet highlights the limitations of existing mathematical models to fully reconstitute the circadian system.

**Figure 2.**
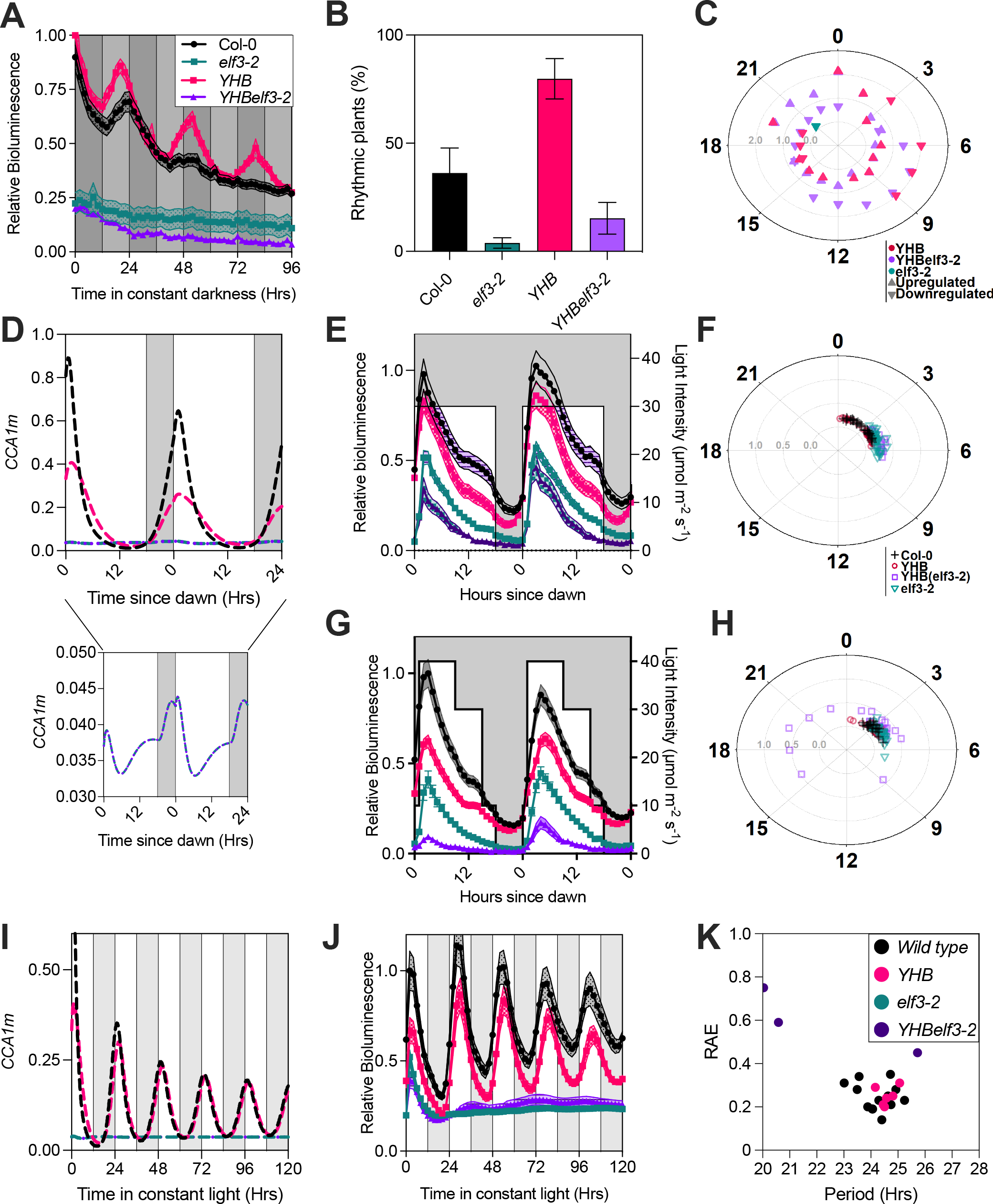
*YHBelf3-2* plants lack circadian rhythms but retain modest responses to light cues. (A) Waveforms of luciferase bioluminescence rhythms of wild-type (Col-0; black), *YHB* (pink), *elf3-2* (green), and *YHBelf3-2* (purple) seedlings expressing a *CCA1::LUC2* reporter, entrained for 7 days under 12 hr:12 hr light:dark cycles (indicated before timepoint 0 by white and grey bars respectively) before transfer to constant darkness (with subjective day:night cycles in constant darkness indicated by grey and light grey bars after timepoint 0). **(B)** Percentage of seedlings measured in (A) which presented robust circadian rhythms [calculated using BioDare; biodare2.ed.ac.uk; (Zielinski et al., 2014)]. Data are presented as mean ± SEM from three independent experiments. **(C)** Phase distribution plot showing phase of differentially expressed genes (log2FC > 1.0 and *p* < 0.05) in 12-day old *YHB, elf3-2*, and *YHBelf3-2* plants relative to Col-0. Plants were harvested 48 hours after transfer to constant darkness. Pyramids indicate up-regulated genes, inverted pyramids represent down-regulated genes. Filled symbols denote significantly enriched phase, colours as in (A). **(D)** Modelled accumulation of *CCA1m* (*CCA1* mRNA) in 12:12 light:dark cycles. Dark grey bars indicate periods of darkness. **(E)** Patterns of luciferase bioluminescence rhythms of Col-0, *YHB*, *YHBelf3-2* and *elf3-2* seedlings expressing a *CCA1::LUC2* reporter in 12:12 light:dark cycles. **(F)** Phase distribution plot showing time of peak *CCA1-*driven luciferase bioluminescence calculated from (D). **(G)** Patterns of luciferase bioluminescence rhythms of Col-0, *YHB*, *YHBelf3-2* and *elf3-2* seedlings expressing a *CCA1::LUC2* reporter, entrained for 7 days in variable light conditions (cycles of 1hrs 10 µmol m^-2^ s^-1^, 8hrs 40 µmol m^-2^ s^-1^, 6hrs 30 µmol m^-2^ s^-1^, 3hrs 10 µmol m^-2^ s^-1^ white light followed by 6hrs of darkness). **(H)** Phase distribution plot showing time of peak *CCA1-*driven luciferase bioluminescence calculated from (G). **(I)** Modelled accumulation of *CCA1m* (*CCA1* mRNA) in constant light. Light grey bars indicate periods of subjective darkness. **(J)** Waveforms of luciferase bioluminescence rhythms of wild type, *elf3-2*, *YHB*, and *YHBelf3-2* seedlings expressing a *CCA1::LUC2* reporter, entrained for 7 days under 12 hr:12 hr light:dark cycles and constant 22 °C temperature before transfer to constant light for imaging **(K)** Assessment of rhythmic robustness (Relative Amplitude Error, RAE) plotted against circadian free-running period for data presented in (J). Experimental data are representative of 3 independent experiments (n ≥15). Error bars indicate SEM.

### ELF3 and YHB have a reciprocal epistatic relationship spanning circadian and photomorphogenic gene expression programs

To further explore the regulation of gene expression in *YHBelf3-2* seedlings we used RNA sequencing to assess transcript accumulation in plants 48hrs after transfer to constant darkness at dusk (ZT60), a time point at which wild-type seedlings appeared to have become arrhythmic and therefore had relatively stable levels of circadian-controlled transcript abundance (Figs. 1C, 2A, S4). Log fold change (-log2FC) in each of *elf3-2*, *YHB,* and *YHBelf3-2* plants was determined relative to wild type (Fig. S4, Table S1). As expected, GO terms associated with responses to light stimuli were over-represented in our lists of genes mis-expressed in *YHB* and *YHBelf3-2* (Fig. S5, Table S2). GO terms associated with circadian rhythms were also significantly over-represented (Fig. S5, Table S2). We next examined whether mis-regulated transcripts in each genotype tended to be expressed at particular times of day [Fig. 2C; (Bonnot et al., 2022)]. Significantly mis-accumulated transcripts were not confined to a single time period in *elf3-2*, *YHB,* or *YHBelf3-2* lines, suggesting that the circadian system is not ‘locked’ at a particular circadian phase in any of these genotypes (Fig. 2C). Instead, differences in the accumulation of numerous core circadian transcripts were apparent [Fig. S4A-D; (Hsu and Harmer, 2014, Laosuntisuk et al., 2023)]. In constant darkness, *elf3-2* plants accumulate increased levels of *GIGANTEA, PRR9,* and *BROTHER OF LUX ARRHYTHMO* whereas *CCA1, LHY,* and *REVEILLE8* (*RVE8*) steady-state levels are reduced (Fig. S4A). 10 of the notional 60 core clock genes are highly mis-accumulated in *YHB* relative to wild type (9 upregulated and 1 downregulated; Fig. S4A, S4C). Interestingly, we mainly observed additive effects in the combined *YHBelf3-2* line (Fig. S4A, S4C-D). 7 of the upregulated genes in *YHBelf3-2* are similarly upregulated in *YHB* while an additional 3 upregulated and 1 downregulated transcript in *YHBelf3-2* were similarly differentially expressed in *elf3-2*. Interestingly, *REVEILLE 8* (*RVE8*) was upregulated in *YHB* (Fig.S4C, p = 6.190x10^-5^) while downregulated in *elf3-2* (Fig. S4B, p = 0.011). *RVE8* was not differentially regulated in *YHBelf3- 2* (Fig. S4C, p-value = 0.829). These data suggest antagonistic regulation of *RVE8* by phyB and ELF3. Our data are generally consistent with previous literature noting that phyB and ELF3 affect transcript accumulation via independent pathways although the necessity of ELF3 to maintain circadian rhythms ensures that *elf3-2* is epistatic to *YHB* in the maintenance of circadian rhythms [Fig. 2A, S4; (Thines and Harmon, 2010)].

To further address how YHB and ELF3 govern photomorphogenesis, we examined differential expression of genes associated with a response to light stimuli [GO:0009416; (Ashburner et al., 2000, Gene et al., 2023)] using our RNAseq dataset of dark-adapted plants (Fig. S4E-H). Of the 740 light stimulus-associated transcripts examined, only 33 are mis-regulated in *elf3-2* plants, with six downregulated and 27 upregulated transcripts (Table S1). Of these, *elf3-2* and *YHBelf3-2* plants share only 7 mis-regulated transcripts, one of which (*HOMEOBOX-LEUCINE ZIPPER PROTEIN 4; HB4*) has previously been shown to play a role in shade avoidance via both phytochrome signalling and ELF3 [Fig. S4E-H, (Sorin et al., 2009, Jiang et al., 2019)]. *HB4* is downregulated in *elf3-2* but upregulated in *YHB* and *YHBelf3-2* (Fig. S4E-H, Table S1). By contrast, 113 transcripts are significantly differentially expressed in *YHB* plants relative to wild type, with 83 of these being upregulated and 30 downregulated (Figs. S4G-H, Table S1). 91 of these transcripts are similarly differentially expressed in both *YHB* and *YHBelf3-2* plants (Fig. S4G-H). These data suggest that disruption of *ELF3* has a modest effect upon YHB-induced light responses, reaffirming that phyB- and ELF3-mediated effects upon light responsive transcripts are genetically separable (Nieto et al., 2022). We did, however, note that two light stimulus responsive genes are similarly differentially expressed in *elf3-2*, *YHB* and *YHBelf3-2*; *RESPONSE REGULATOR 3* (*ARR3*), which is strongly downregulated in all three genotypes and *COLD REGULATED GENE 27* (*COR27*) which is strongly upregulated (Table S1). Both genes have been previously shown to interact with the circadian clock (Salome et al., 2006, Zhu et al., 2020). Our data illustrate the epistatic role of phyB signalling upon photomorphogenesis despite the inter-related nature of phyB- and ELF3-mediated effects upon gene expression.

### *YHBelf3* plants have a reduced response to fluctuating environmental conditions

We next examined the behaviour of *YHBelf3* seedlings in the presence of light. Although our modelling expected that *elf3* and *YHBelf3* would essentially be arhythmic in response to dawn and dusk (Fig. 2D), each of the genotypes examined displayed circadian entrainment to experimental light signals and retained daily responses to dawn, as measured by the expression of *CCA1::LUC2* bioluminescence in driven light:dark cycles (Fig. 2E-F). *CCA1::LUC2* bioluminescence begin to increase in wild-type and *YHB* seedlings 1-3 hours before dawn, indicating a circadian anticipation of dawn in these plants (Fig. 2E). By contrast, this dawn anticipation was absent in *elf3-2* and *YHBelf3-2* plants, with *CCA1::LUC2* driven bioluminescence only beginning to increase after the application of light (Fig. 2E).

Since *elf3-2* and *YHBelf3* plants retained a response to dawn, we examined the activation of the *CCA1* promoter in response to varied light intensity during the photoperiod (Fig. 2G-H). A ‘variable light’ regime was designed, where light intensity varied in a pseudo-sinusoidal pattern throughout the day, peaking in the late morning and gradually decreasing as dusk approached (Fig. 2G). Our experimental data demonstrated that *elf3-2* retained entrainment to variable lighting, although *YHBelf3-2* was unable to entrain to these conditions (Fig. 2G-H). These data are consistent with additional photosensory systems feeding into the regulation of *CCA1*, including metabolic signals from photosynthesis (Jones, 2018, Jones, 2019, Wang et al., 2024, Haydon et al., 2013).

We next assessed circadian rhythmicity in *YHBelf3* seedlings held in constant light. Our modelling predicted that wild-type and *YHB* seedlings would have comparable circadian rhythms in constant light (Fig. 2I). In line with this hypothesis, circadian rhythms in *YHB* seedlings were indistinguishable from wild type in constant white light (Fig. 2J-K). Similarly, circadian rhythms dissipated in *elf3-2* and *YHBelf3-2* seedlings within 24 hours of transfer to constant white light (Fig. 2J-K).

### *YHBelf3-2* have reduced growth and flowering plasticity in response to light and temperature cues

The combination of *YHB* and *elf3* alleles decouples the circadian system from photomorphogenesis, although *YHBelf3-2* plants can retain daily patterns of gene expression when grown in light:dark cycles (Figs. 1 + 2). We were therefore interested how our genetic manipulations affected developmental traits and life cycle transitions in varied light conditions (Fig. 3). Our FMv2+F2014 model predicted that hypocotyl length would be uncoupled from photoperiod (Fig. 3A). We observed that *YHB*-driven growth phenotypes persisted in the hypocotyls of 5-day old seedlings, with *YHBelf3-2* seedlings retaining a short hypocotyl phenotype regardless of the light condition utilised for growth (Fig. 3B-D). We note that *YHB* and *YHBelf3- 2* seedlings were indistinguishable from wild type when grown under long day conditions (Fig. 3D).

**Figure 3.**
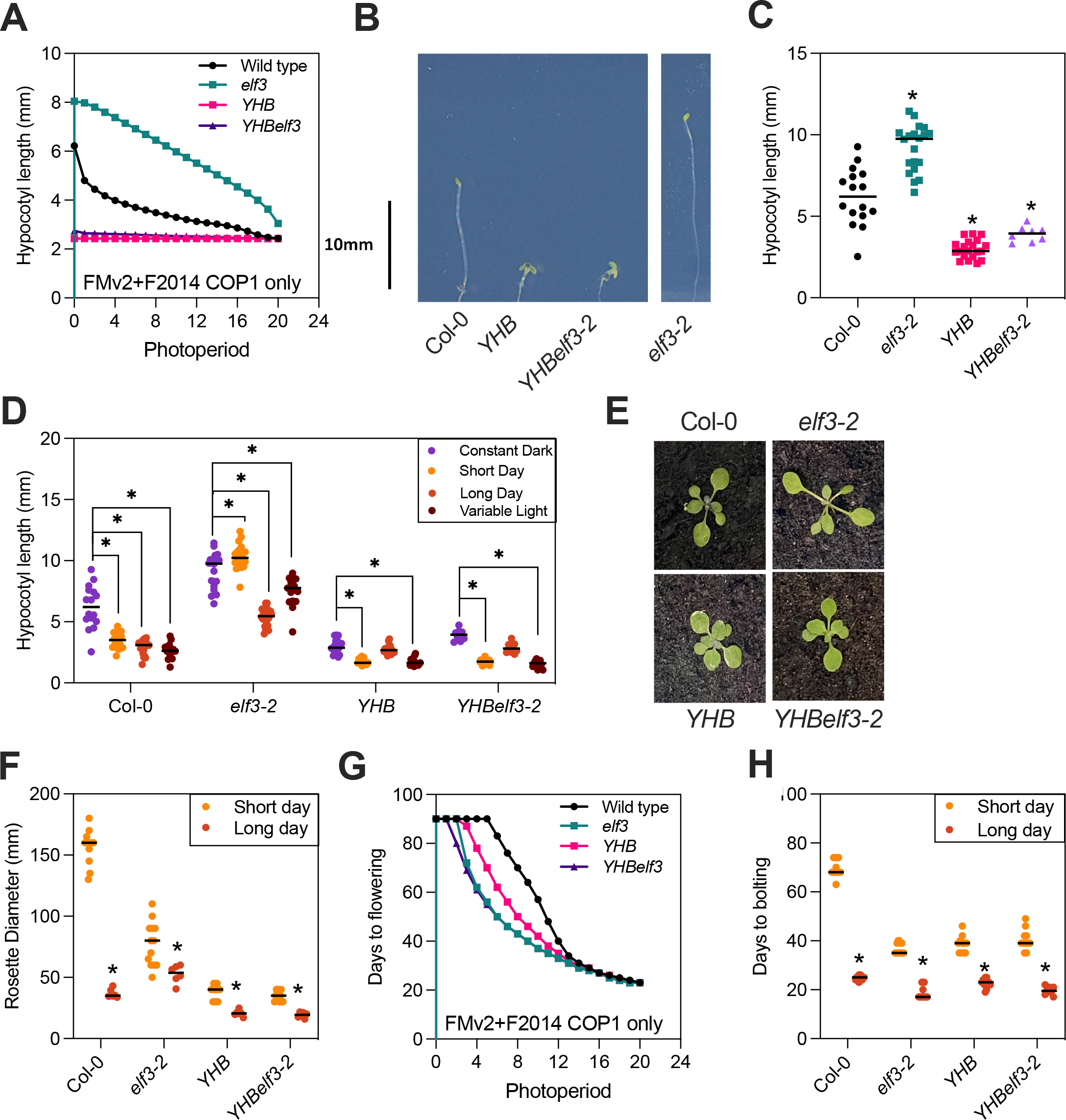
*YHBelf3* plants are less responsive to changing light environments. (A) Modelled hypocotyl length in wild type, *elf3*, *YHB*, and *YHBelf3* seedlings under different simulated photoperiods. **(B)** Representative images of wild type (WT; Col-0), *YHB*, *YHBelf3-2* and *elf3-2* seedlings grown vertically on 0.5 MS plates for five days in constant darkness. **(C)** Quantification of the hypocotyl lengths of Col-0, *YHB*, *YHBelf3-2* and *elf3-2* seedlings grown vertically on 0.5 MS plates for five days in constant darkness. Data shows a representative example from 3 independent experiments (n ≥9). **(D)** Hypocotyl length of Col-0, *elf3-2*, *YHB* and *YHBelf3-2* seedlings grown vertically on 0.5 MS plates for five days in constant darkness (purple), short day cycles (yellow), long day cycles (orange) or variable-light cycles (brown; cycles of 1hrs 10 µmol m^-2^ s^-1^, 8hrs 40 µmol m^-2^ s^-1^, 6hrs 30 µmol m^-2^ s^-1^, 3hrs 10 µmol m^-2^ s^-1^ white light followed by 6hrs of darkness). **(E)** Representative images of wild type (Col-0), *YHB*, *YHBelf3-2* and *elf3-2* seedlings grown on soil for 21 days under long day cycles (18 hr:16 hr light:dark) with 150 µmol m^-2^ s^-1^ white light and a constant temperature of 22 °C. **(F)** Rosette diameter of 28 day old Col-0, *elf3-2*, *YHB* and *YHBelf3-2* seedlings grown on soil under short or long days at 22 °C. **(G)** Modelled flowering time in wild type, *elf3*, *YHB*, and *YHBelf3* seedlings under different simulated photoperiods. **(H)** Flowering time of wild type, *YHB*, *YHBelf3-2* and *elf3-2* plants grown on soil at a constant temperature of 22 °C under long- or short- days. Data shows a representative example from 3 independent experiments (n ≥10).

We next examined growth phenotypes in more mature Arabidopsis plants (3 weeks after germination; Fig. 3E-F, S6). The size of wild-type Arabidopsis plants is greatly dependent upon photoperiod length when plants are grown at 22°C, with rosette diameter decreasing as photoperiod increases (Fig. 3E-F). Regardless of day length, *elf3-2* seedlings had an expanded rosette diameter compared to wild-types grown under long days, possibly related to the loss of light perception in these lines [Fig. 3E-F; (Zagotta et al., 1996)]. We noted substantial variation in rosette diameter in wild-type and *elf3-2* plants, although rosette diameter was more consistent under longer photoperiods (Fig. 3E-F). By contrast, the rosette of *YHB* and *YHBelf3-2* seedlings was more compact and uniform in size regardless of daylength (Fig. 3E-F).

*YHB* and *elf3-2* genotypes have both previously been shown to have an early flowering phenotype when grown under short-day conditions and so we expected that this phenotype would be shared by *YHBelf3-2* plants [Fig. 3G-H; (Franklin and Quail, 2010, Hajdu et al., 2015, Zagotta et al., 1996)]. Our FMv2+F2014 model similarly predicts that *YHBelf3* plants will display reduced photoperiodic sensitivity comparable to *elf3* (Fig. 3G). In agreement with this hypothesis, *YHB*, *elf3*, and *YHBelf3-2* plants flowered earlier than wild type under either long-days or short-day conditions (Fig. 3G-H).

Both phyB and ELF3 have been shown to be critical for temperature responses in addition to their roles in photoperception (Jung et al., 2020, Jung et al., 2016, Legris et al., 2016). We therefore compared how our *YHBelf3* plants performed under varied temperature conditions (Fig. 4). In contrast to light-driven entrainment (Figs. 2E-F), *CCA1*-driven bioluminescence peaked 6 hours after dawn in wild-type when entrained to temperature (Fig. 4A-B). The phase of *CCA1-*driven bioluminescence was unaffected in *YHB* seedlings, although neither *elf3-2* nor *YHBelf3* seedlings were able to entrain to temperature signals when held in constant light (Fig. 4A-B). These data suggest that light cues are necessary to drive rhythmic *CCA1* expression in *elf3* and *YHBelf3* seedlings.

**Figure 4.**
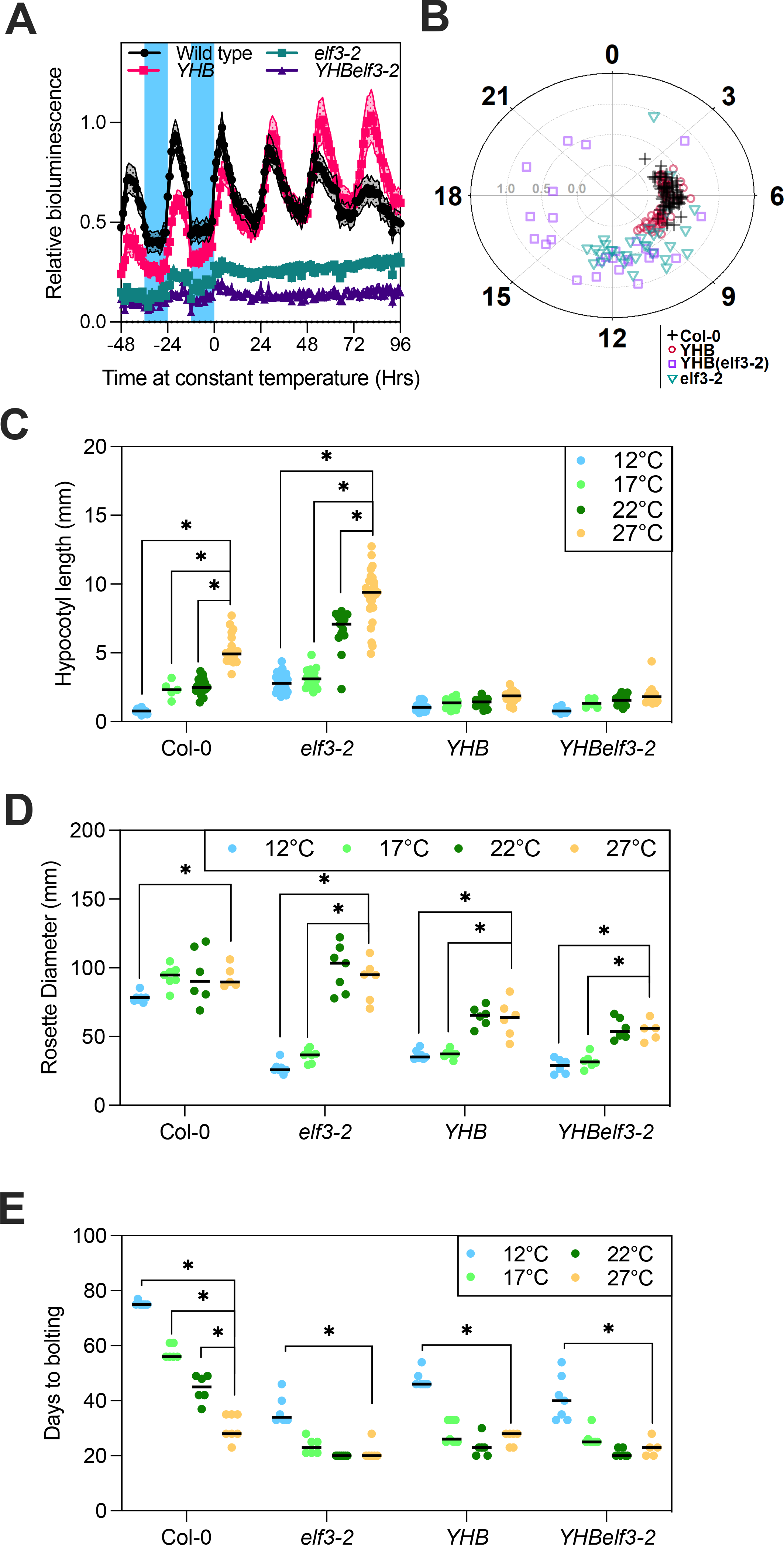
*YHBelf3* plants are less responsive to temperature-driven environmental cues. (A) Patterns of luciferase bioluminescence rhythms of wild type, *elf3-2*, *YHB*, and *YHBelf3-2* seedlings expressing a *CCA1::LUC2* reporter, entrained for 7 days under 12 hr:12 hr 22 °C:17°C cycles and constant white light before transfer to testing conditions at a constant temperature of 22°C. **(B)** Phase distribution plot showing time of peak *CCA1-*driven luciferase bioluminescence calculated from (A). Data are presented as the mean ± SEM and are representative of at least 3 independent experiments (n ≥ 15). **(C)** Hypocotyl length of Col-0, *elf3-2*, *YHB* and *YHBelf3-2* seedlings grown vertically on 0.5 MS plates for five days under 12 hr:12 hr light:dark cycles at a constant temperature of (from left to right) 12 °C (blue), 17 °C (light green), 22 °C (dark green), or 27 °C (yellow). **(D)** Rosette diameter of 28 day old Col-0, *elf3-2*, *YHB* and *YHBelf3-2* seedlings grown on soil under 12 hr:12 hr light:dark cycles at a constant temperature of (from left to right) 12 °C (blue), 17 °C (light green), 22 °C (dark green) or 27 °C (yellow). **(E)** Flowering time of wild type, *YHB*, *YHBelf3-2* and *elf3-2* plants grown on soil under 12 hr:12 hr light:dark cycles at a constant temperature of (from left to right) 12 °C (blue), 17 °C (light green), 22 °C (dark green) or 27 °C (yellow). Data are representative of at least three biological repeats. Error bars indicate SEM. See also Figure S6.

As under varied light, *YHBelf3-2* hypocotyl growth was unaffected by temperature, with no significant difference in hypocotyl extension between 12°C and 27°C (Fig. 4C). Seedling growth is therefore more uniform in *YHBelf3-2* plants regardless of light or temperature cues. Ambient temperature also affected rosette diameter (Figs. 4D, S6). Wild-type plants maintain a comparatively consistent diameter between 12°C and 27°C when grown in neutral day conditions (12:12 light:dark cycles), with a modest yet significant decrease at 12°C (Fig. 4D, Fig. S6). By contrast, *elf3-2* seedlings were more sensitive to lower temperatures, with rosette diameter being substantially decreased at 12°C and 17°C compared to higher temperatures (Fig. 4D). *YHB* and *YHBelf3-2* plants were also responsive to these temperature changes although the difference in size was smaller than observed in *elf3-2* plants (Fig. 4D). Under neutral day conditions, flowering was delayed in all genotypes when plants were grown at 12°C, with flowering time accelerating in wild-type plants as temperatures increased (Fig. 4E). By contrast, *YHB*, *elf3,* and *YHBelf3* genotypes retained stable flowering times from 17°C to 27°C (Fig. 4E). The *YHB* and *YHBelf3* plants therefore retain uniform and early flowering phenotypes and so demonstrate reduced developmental plasticity across a range of light conditions and temperatures.

### Reducing developmental plasticity has potential beneficial Implications for farming

Photo- and thermo-morphogenesis are crucial processes that enable plants to optimise growth and development in response to prevailing environmental conditions by developmental plasticity. Our data validate mathematical models and demonstrate that expression of *YHB* is epistatic to the morphological consequences of *ELF3* disruption, although *ELF3* is essential to maintain circadian rhythmicity (Fig.1-3). *YHBelf3-2* plants consequently retain a vegetative phenotype comparable to wild-type yet have an early flowering phenotype and are unable to anticipate environmental transitions (Figs. 3-4). The combination of *YHB* and *elf3* alleles consequently produces plants less responsive to environmental signals that retain vegetative growth and predictable flowering. This demonstrates how engineering of the circadian system alongside environmental signalling pathways creates plants with reduced developmental plasticity.

Although developmental plasticity is advantageous in natural conditions (where competition for resources and environmental stresses vary across seasons and location) this trait is disadvantageous in modern crop monoculture where fertilisers, pesticides, irrigation, etc., can be provided. One goal of modern breeding programmes is to increase the uniformity of crops so that harvesting time is more predictable and quality is consistent. This is true for intensive, precision outdoor farming and Total Controlled Environment Agriculture (TCEA, or vertical farming). In addition, climate change has rapidly altered daylength and temperature relationships worldwide, and maintaining crop productivity in current locations or moving to more favourable temperate latitudes will require manipulation of environmental responses. Our modelling predicted that manipulation of phyB and ELF3 signalling cascades would enable the restriction of developmental plasticity in plants to changing photoperiods (Figs. 1-3). Crucially, we have been able so show that the combination of these two alleles [*YHBelf3*] has advantages over each allele individually. *YHBelf3- 2* plants retain consistently earlier flowering times without impairing vegetative growth (Fig. 3-4). Since ELF3 and YHB have conserved function across species it will be of great interest to apply these genetic modifications to reduce developmental variation in crop species (Huang et al., 2017, Hu et al., 2020).

## Methods

### Plant Material and Growth Conditions

Wild type *CCA1::LUC+*, and *elf3-2 CCA1::LUC+* Arabidopsis seed have previously been reported (Huang et al., 2016a). *PHYB::YHB* and *PHYB::YHB* (*elf3-2*) Arabidopsis were generated by transforming *CCA1::LUC+*, and *elf3-2 CCA1::LUC+* seed with pJM63 gYHB (Su and Lagarias, 2007) via floral dip (Clough and Bent, 1998). Transformants were selected with 75μg mL^-1^ kanamycin to identify homozygous seedlings in the T3 generation. *phyb-9 elf3-1* lines were generated by crossing *elf3-1* to CCA1::LUC+ and *phyB-9* was crossed to CCA1::LUC+, with long hypocotyl, bioluminescent F2 seedlings confirmed for homozygous *elf3-1* and *phyB-9* alleles using a dCAPS primer strategy as described previously (Nusinow et al., 2011, Huang et al., 2016b). *elf3-1* CCA1::LUC+ was then crossed to *elf3-1 phyB-9* (Reed et al., 2000), and bioluminescent, long hypocotyl F2 lines were confirmed as *elf3-1 phyB-9* using dCAPS primers. F3 lines were screened for bioluminescence to identify homozygous CCA1::LUC+ seedlings.

CER was cloned from plasmid CER C1 (Koushik et al., 2006) using primers pDAN0869 and pDAN0870 and recombined with pB7-SHHc (Huang et al., 2016b) digested with AvrII using In- Fusion HD cloning (Clontech, Mountain View, CA) to generate pB7-CER-SHHc. pENTR-YHB (Huang et al., 2016b) was recombined with pB7-CER-SHHc to generate pB7-YHB-CER-SHHc. This plasmid was transformed into *elf3-1 phyb-9 CCA1::LUC+* to generate *35S::PHYB(elf3-1 phyb-*9) *CCA1::LUC+* and transformants were identified by BASTA resistance.

T3 and F3 seed were surface sterilised in chlorine gas prior to plating on half-strength Murashige and Skoog (0.5 MS) medium (Prasetyaningrum et al., 2023). Seedlings were entrained for 5–12 d before being transferred to testing conditions as described in each figure legend. During standard growth, plants were kept under 150 µmol m^−2^ s^−1^ white light in 12 hrs:12 hrs light:dark cycles in Panasonic MLR-352-PE chambers. Relative humidity and temperature were set to 60– 70% and 22°C, respectively except where growth under other temperatures conditions are listed.

### Hypocotyl assays

Seeds were grown on 0.5 MS agar plates and irradiated with cool fluorescent white light at 170 μmol m^−2^ s^−1^ for 4 hr before being moved to LED chambers as per experimental requirements and grown vertically for 5 days before being imaged and processed using ImageJ (Schneider et al., 2012). Short day, long day and squareform treatments used 30 μmol m^−2^ s^−1^ and the fluctuating light treatment used a cycle of 1hr 10 μmol m^−2^ s^−1^, 8hrs 40 μmol m^−2^ s^−1^, 5hrs 30 μmol m^−2^ s^−1^, 4hrs 10 μmol m^−2^ s^−1^ and 6hrs darkness. Data were plotted and analysed using a One-way ANOVA followed by Dunnett’s multiple comparisons test in GraphPad Prism version 10.0.3.

### Luciferase assays

Individual seedlings were grown for 6 days in 12:12 light:dark cycles under white light on half-strength MS media as in previous work (Prasetyaningrum et al., 2023). Plants were sprayed with 3 mM D-luciferin in 0.1% Triton X-100, before being transferred to imaging conditions as described for each experiment. Individual plants were imaged repeatedly (every 1-2 hours) dependent upon the experiment using a Retiga LUMO camera run by MicroManager 1.4.23(Edelstein et al., 2014) using a custom script. In experiments where temperature was not constant throughout growth and imaging, temperature change was initiated as indicated. The patterns of the luciferase signal were fitted to cosine waves using Fourier Fast Transform-Non-Linear Least Squares [FFT-NLLS; (Plautz et al., 1997)] to estimate the circadian period length made using BioDare2 ((Zielinski et al., 2014); biodare2.ed.ac.uk). Waveforms, periods and percentage rhythmicity data were plotted using GraphPad Prism version 10.0.3.

### qRT-PCR

Following entrainment, seedlings were transferred to constant darkness at dusk. Tissue was harvested and snap-frozen in liquid nitrogen at the indicated time points before RNA extraction using using Tri Reagent® according to the manufacturer’s protocol (Sigma Aldrich, Dorset, UK, http://www.sigmaaldrich.com). Reverse transcription was performed using either Superscript™ II or M-MLV reverse transcriptase according to manufacturer’s protocols (Invitrogen, Waltham, Massachusetts, USA, https://www.thermofisher.com/Invitrogen). Real-time reverse transcription polymerase chain reaction was performed using a QuantStudio™ 3 Real-Time PCR System or a StepOnePlus™ Real-Time PCR System (Applied Biosystems, Waltham, Massachusetts, USA, https://www.thermofisher.com/AppliedBiosystems). Samples were run in triplicate and starting quantities were estimated from a critical threshold using the standard curve of amplification.

*APA1*, *APX3* and *IPP2* expression was used as an internal control, with data for each sample normalised to these as previously described (Nusinow et al., 2011).

### RNAseq

Plants were grown on 0.5 MS agar plates under entrainment conditions for 12 days. At dusk on the twelfth day of growth (ZT12), seedlings were transferred to constant darkness. Pools of ca. 20 seedlings were harvested and snap-frozen in liquid nitrogen 48 hours later (ZT60). Total RNA was extracted from three biological replicates per genotype using Tri Reagent® according to the manufacturer’s protocol (Sigma Aldrich, Dorset, UK, http://www.sigmaaldrich.com). Library preparation and Illumina sequencing (Illumina, San Diego, USA) with 150bp paired-end reads was performed by Novogene Biotech (Cambridge, UK) using Illumina protocols. RNAseq reads were first aligned and to the AtRTD3 transcriptome (Zhang et al., 2022) and read-counts were generated using Kallisto (Bray et al., 2016) in the Galaxy platform (Afgan et al., 2016).

Subsequent analysis was performed using the 3DRNAseq pipeline (Guo et al., 2021). Normalised transcript abundance in wild type samples was used to calculate fold change difference of transcript abundance. A list of transcripts contributing to circadian rhythmicity were derived from (Hsu and Harmer, 2014, Laosuntisuk et al., 2023). Gene ontology annotation was performed using DAVID (Huang et al., 2009, Sherman et al., 2022). A list of 740 genes were taken from the GO term GO:0009416 response to light stimulus (Ashburner et al., 2000, Gene et al., 2023) https://www.arabidopsis.org/servlets/TairObject?type=keyword&id=6751]. Genes of interest were plotted in heatmaps and volcano plots using R (R Core Team, 2013) and RStudio (Posit Software).

### Flowering time and growth analysis

Following stratification, plants were grown on soil until bolting. Rosette area, rosette diameter and leaf counts were measured regularly throughout the growth period (ca. twice per week). The number of days to bolting were recorded when the bolt was 1cm above the rosette. Plants were grown under 150 µmol m^-2^ s^-1^ white light with day length and temperature varied between experiments. For variable day length experiments, plants were grown under long-days (16 hrs light: 8 hrs darkness) or short-days (8 hrs light: 16 hrs darkness) at 22°C. For temperature response experiments, plants were grown under balanced day lengths (12 hrs light: 12h hrs dark) under either 27°C, 22°C, 17°C or 12°C. Data were plotted and analysed using a two-way ANOVA followed by Tukey’s multiple comparisons test in GraphPad Prism version 10.0.3.

### Mathematical modelling

The Arabidopsis Framework Model version 2 (FMv2)(Chew et al., 2022) is a multiscale model of Arabidopsis that brings together multiple modules to describe diverse processes including the circadian clock, flowering, metabolism and vegetative growth. The F2014 model (Fogelmark and Troein, 2014) is an updated Arabidopsis circadian clock model with fewer explicit light-sensitive reactions and without extended transcriptional activation. Both these models were used and combined in this study. The original FMv2 model was simulated, with minimal changes as described below to allow for introduction of the YHB mutant and for model comparison. The “FMv2+F2014” model was constructed by replacing the P2011 (Pokhilko et al., 2012) circadian module of FMv2 with the updated F2014 circadian model, in the spirit of the modular Framework model.

### FMv2 model

The MATLAB code for the FMv2 was downloaded from the github repository: https://github.com/danielseaton/frameworkmodel/ (FAIRDOM link: https://fairdomhub.org/models/248) and run in MATLAB R2022a.

### Addition of F2014

MATLAB code was written to simulate the F2014 model based on the equations described in (Fogelmark and Troein, 2014). ChatGPT4 was initially used to convert the PDF image of the equations into LaTeX code. This was then manually corrected to remove errors introduced by the AI and then converted from LaTeX into MATLAB manually.

Conversion to MATLAB was also performed using ChatGPT4, and the two were compared as an additional check.

The F2014 model replaced the P2011 module of the FMv2 model. Scaling factors were added to rescale the amplitudes of the outputs of the circadian module F2014 to match those of P2011, to allow input to the PIF-CO-FT (Seaton-Smith) module (Seaton et al., 2015). Furthermore, CCA1 and LHY are modelled separately in F2014, so the sum of the two was used to replace the LHY input to the PIF-CO-FT module. Specifically:

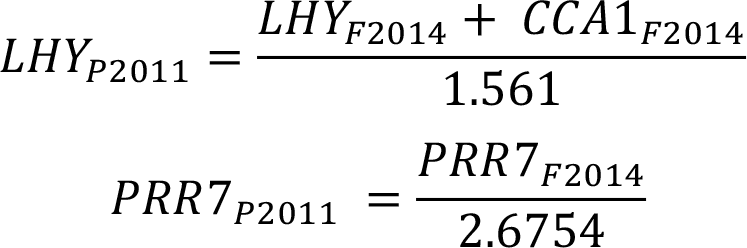

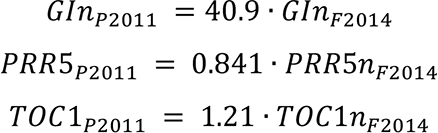

### Parameter choice

The parameter set 1 of (Fogelmark and Troein, 2014) was used in all simulations of this model. Parameters as preset in FMv2 were used for all other modules with the exception of parameters for the hypocotyl length calculation and the photothermal time threshold for flowering, which were calibrated for the laboratory’s wildtype data for various photoperiods (Fig. S1b). These parameters were used unchanged for the mutant predictions.

Flowering Threshold: A single parameter value was used for both the FMv2 and the FMv2+F2014 models. The threshold value was 4107.6 MPTU.

The Hypocotyl length was calculated according to the equation used in (Seaton et al., 2015) :

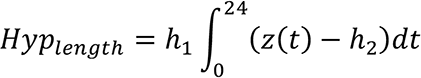

Where

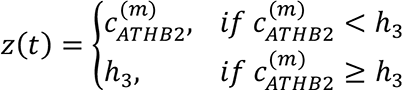

Reparameterisation was carried out for *h*_1_, *h*_2_, *h*_3_ separately for each version of the model.

### Simulating Mutants

The *elf3* and YHB mutations were introduced in both P2011 and F2014 models. The *elf3* mutation is present in the original code for FMv2 (P2011), so this was simulated in the same way. For F2014, the ELF3 protein production parameter p16 was set to 0 in the mutant. The YHB mutant was added in both models, either “Globally” by altering all light inputs except for blue light (assumed to affect the GI and ZTL protein light-sensitivities and the dark accumulator) or by altering only COP1-related light inputs. The alteration in both cases was to set the light to be 75% ON in the dark, to account for the constitutively active phyB signalling.

YHB is also affecting the PIF-CO-FT module directly, where phyB is explicitly modelled. In this case, the light variable for the phyB equation is set to 1 at all times in the mutant.

### Model simulation

The ODEs were solved numerically using MATLAB’s ode15s. The circadian module for both P2011 and F2014 was initialised and entrained for 12 days in 12L/12d conditions prior to the simulation start. Initial conditions were set as in (Chew et al., 2022) for P2011, while for F2014 the initial value 0.1 was used for all variables.

### Data Availability

Further information and requests for resources and reagents should be directed to and will be fulfilled by Matt Jones. Plasmids generated in this study are available upon request. RNA-seq data have been deposited at GEO and are publicly available as of the date of publication; GEO: PRJNA1078346. Luciferase data has been deposited in BioDare2 (www.biodare2.ac.uk) and will be publicly available as of the date of publication; accession numbers will be provided here. Any additional information required to re-analyze the data reported in this paper is available from the corresponding authors upon request. Models of hypocotyl growth (Seaton et al., 2015) and flowering time (Chew et al., 2022) are derived from previously published work now available at FAIRDOMHub: https://fairdomhub.org/models/248. All original code will be publicly available as of the date of publication.

## Author contributions

Conceptualization, DAN, MAJ; Methodology, MWB, RAK, DAN, MAJ; Software, RAK; Validation, CD, JO, MWB; Formal Analysis, MWB, SFE, MAJ, JO; Investigation, MWB, SFE, CD, JO; Resources, CD, KNE, RB, MAJ; Data Curation, MWB, SFE, CD, JO; Writing – Original Draft, MWB, MAJ; Visualisation, MWB, SFE; Supervision, DAN, MAJ; Project Administration, DAN, MAJ; Funding Acquisition, DAN, MAJ.

## Acknowledgments

This work was supported by the UKRI (BB/S005404/1, BB/Z514469/1), the Gatsby Charitable Foundation, the Perry Foundation (to SFE), the Douglas Bomford Trust (to SFE) and a William H. Danforth Plant Sciences Fellowship (to KNE).

---

**Figure S1.**
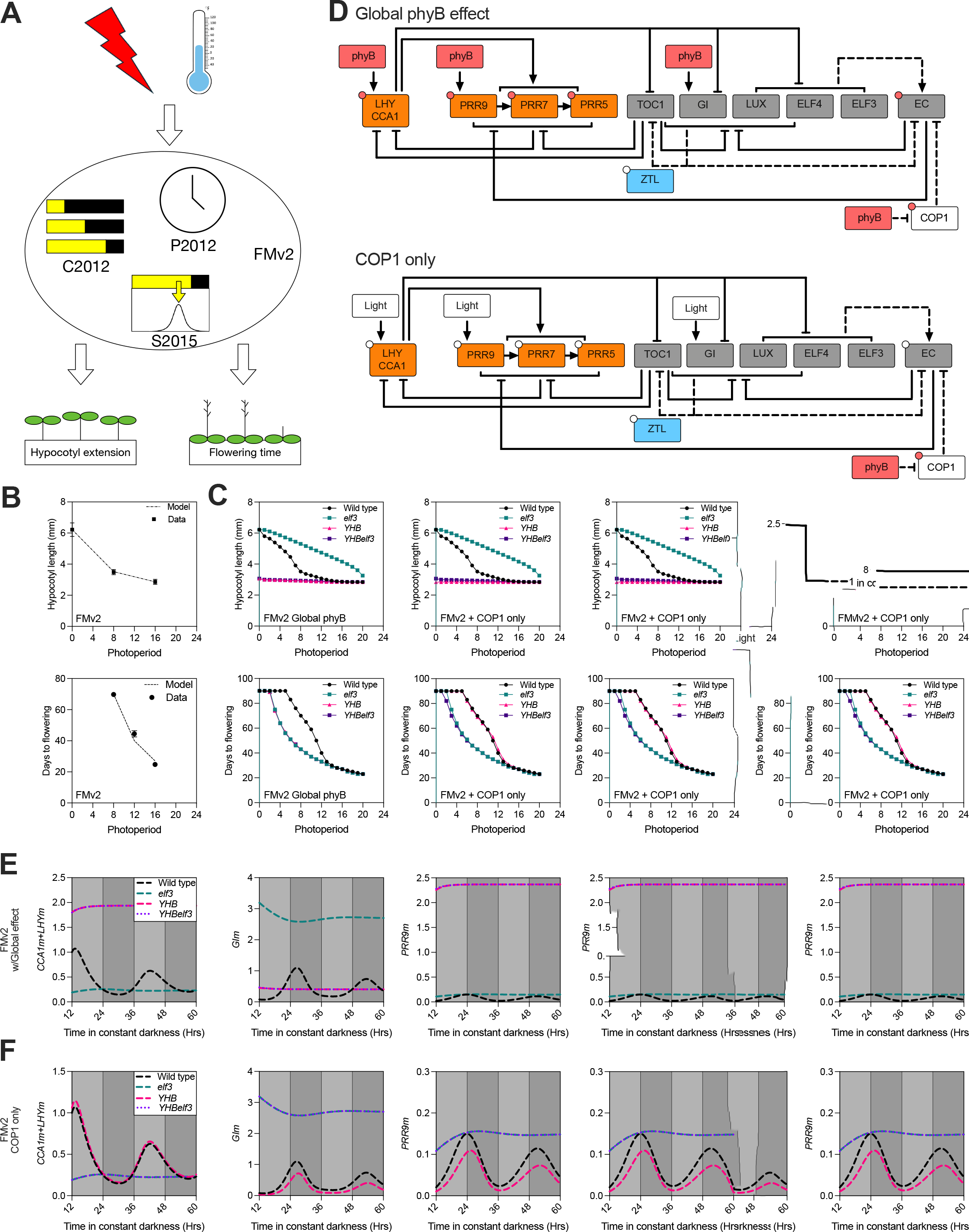
Manipulation of phyB and ELF3 activity is expected to reduce developmental plasticity in plants (A) Cartoon of the revised Arabidopsis Framework Model v2 (FMv2) (Chew et al., 2022). **(B)** Model fitting to hypocotyl length and flowering time. **(C)** Modelled hypocotyl length and flowering time in wild type, *elf3*, *YHB*, and *YHBelf3* seedlings under different simulated photoperiods using either ‘global phyB’ or ‘COP1 only’ FMV2 variants. **(D)** PhyB signalling into the circadian system was modelled via two hypotheses. The ‘Global phyB effect’ variant (upper) proposes that activated phyB is sufficient to induce light-activated gene expression in the circadian system in addition to enabling degradation of COP1. The ‘COP1 only’ variant (lower) restricts the effect of phyB activation solely to the turnover of COP1. In both cases, stability of ZTL and GI is regulated independently since this is a blue light-mediated effect (Kim et al., 2013). Circadian model adapted from P2012 (Pokhilko et al., 2012). Post-translational regulation by light is shown by small white circles. Small red circles indicate post-translational regulation induced by phyB. **(E)** Modelled accumulation of *LHYm* (*CCA1/LHY* mRNA), *GIm* (*GIGANTEA* mRNA), and *PRR9m* (*PRR9* mRNA) in constant darkness following two days of simulated entrainment to 12:12 light:dark cycles using the ‘global phyB’ FMv2 variant. Light grey bars demonstrate subjective day in constant darkness. **(F)** Modelled accumulation of *LHYm* (*CCA1/LHY* mRNA), *GIm* (*GIGANTEA* mRNA), and *PRR9m* (*PRR9* mRNA) in constant darkness following two days of simulated entrainment to 12:12 light:dark cycles using the ‘COP1 only’ FMv2 variant. Light grey bars demonstrate subjective day in constant darkness.

**Figure S2.**
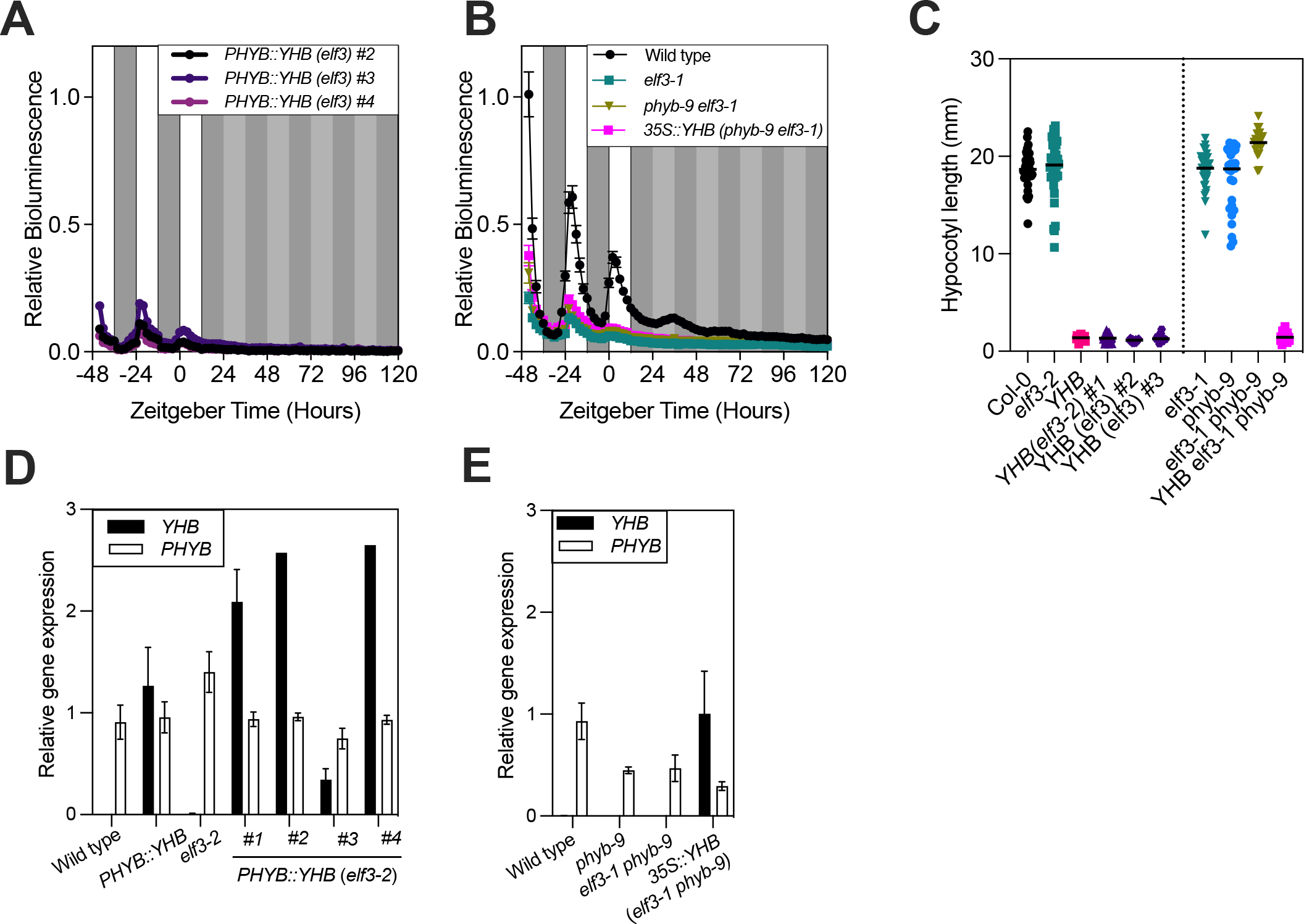
Phenotypes of plants expressing *YHB* under the control of *PHYB* or *35S* promoter in an *elf3* background. (A) Waveforms of luciferase bioluminescence rhythms driven by the *CCA1::LUC+* reporter in independent *YHBelf3-2* lines. Plants were entrained for 7 days under 12 hr:12 hr light:dark cycles (indicated before timepoint 0 by white and grey bars respectively) before transfer to constant darkness (with subjective day:night cycles in constant darkness indicated by grey and light grey bars after timepoint 0). **(B)** Waveforms of luciferase bioluminescence rhythms driven by the *CCA1::LUC+* reporter in Wild-type, *elf3-1*, *phyb-9 elf3-1,* and *35S::YHB phyb-9 elf3-1* seedlings. Plants were entrained and imaged as described in (A). **(C)** Quantification of hypocotyl length in independent lines expressing *YHB* in an *elf3* background. Seedlings were grown vertically on 0.5 MS plates for five days in constant darkness. Data shows a representative example from 3 independent experiments (n ≥8). **(D-E)** qRT-PCR to compare accumulation of endogenous *PHYB* and the *YHB* transgene in the lines examined in this study. Tissue was harvested at ZT2 after 10 days of entrainment. Data is presented relative to *APA1*, *APX3*, and *IPP2*. Error bars indicate SEM. Primers used are listed in Table S4.

**Figure S3.**
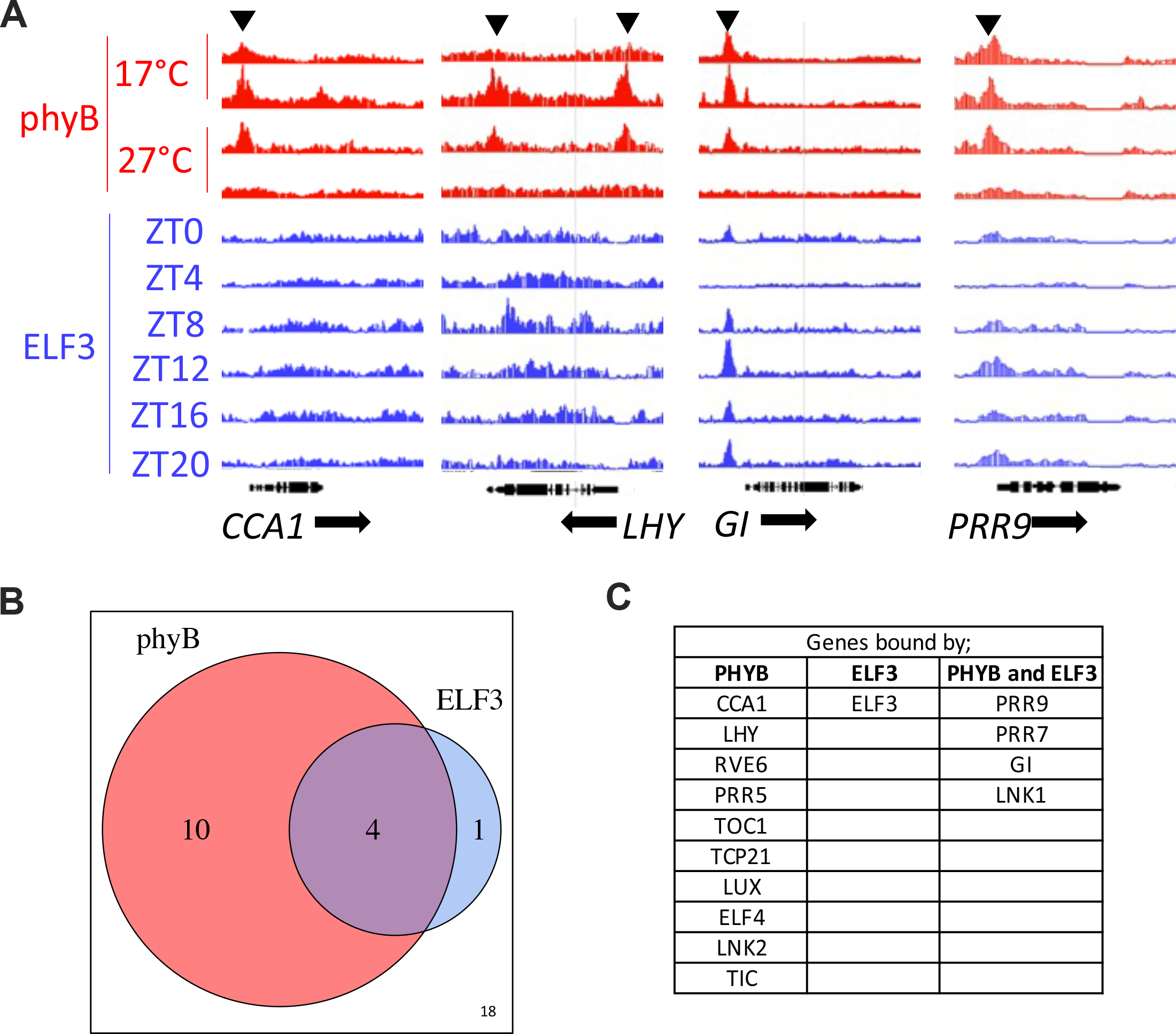
Comparison of ELF3 and PHYB chromatin affinity. (A) Association of phyB and ELF3 with genomic regions encoding selected transcripts. Black arrows indicate significant regions of enrichment indicating phyB (red) or ELF3 (blue) association. Cartoons of gene structure depicting exons and introns are indicated below, along with direction of transcription. **(B)** Venn diagram of circadian genes bound by phyB or ELF3. **(C)** Identities of genes shown in (B). ChIP-seq data is taken from (Jung et al., 2016, Ezer et al., 2017). See also Table S3.

**Figure S4.**
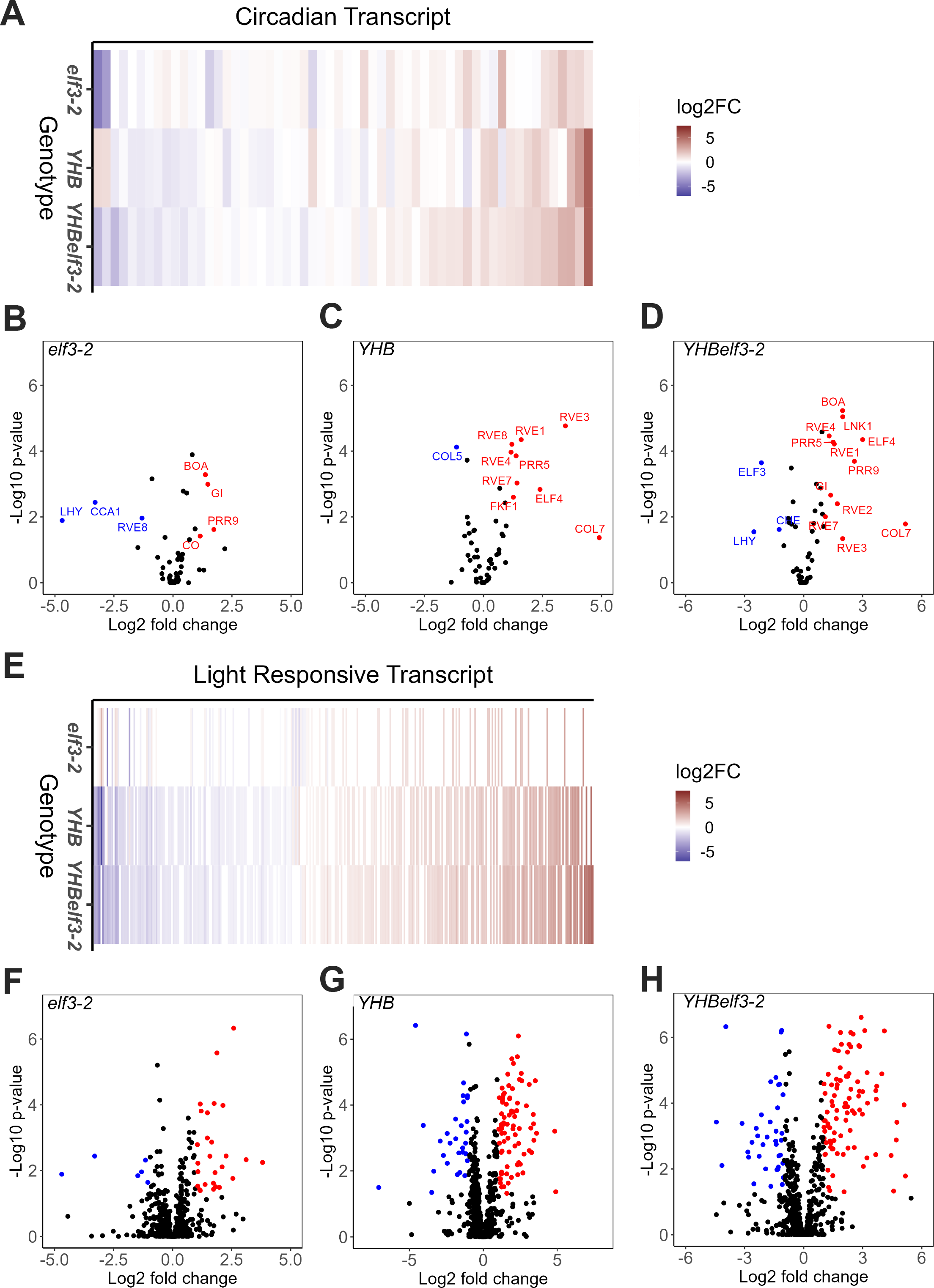
Global differences in transcript accumulation in *YHBelf3* seedlings held in constant darkness. (A) Heat map of significantly differentially accumulated *Arabidopsis* core circadian clock transcripts (*p* < 0.05) in dark-treated *YHB*, *YHBelf3-2* and/or *elf3-2* seedlings relative to dark-treated wild type (Col-0) seedlings. White datapoints indicate no significant difference in expression relative to Col-0 (*p* > 0.05). **(B-D)** Volcano plots showing accumulation of *Arabidopsis* core circadian clock transcripts [list combined from (Hsu and Harmer, 2014, Laosuntisuk et al., 2023, Gene et al., 2023)) in 12-day old seedlings dark-treated for 48 hours prior to harvesting. Data represents transcript abundance in **(F)** *elf3-2*, **(G)** *elf3-2* and **(H)** *YHBelf3-2* relative to wild type (Col-0). Data are plotted using criteria described in (B-D). Data shows the average of the three independent biological replicates.

**Figure S5.**
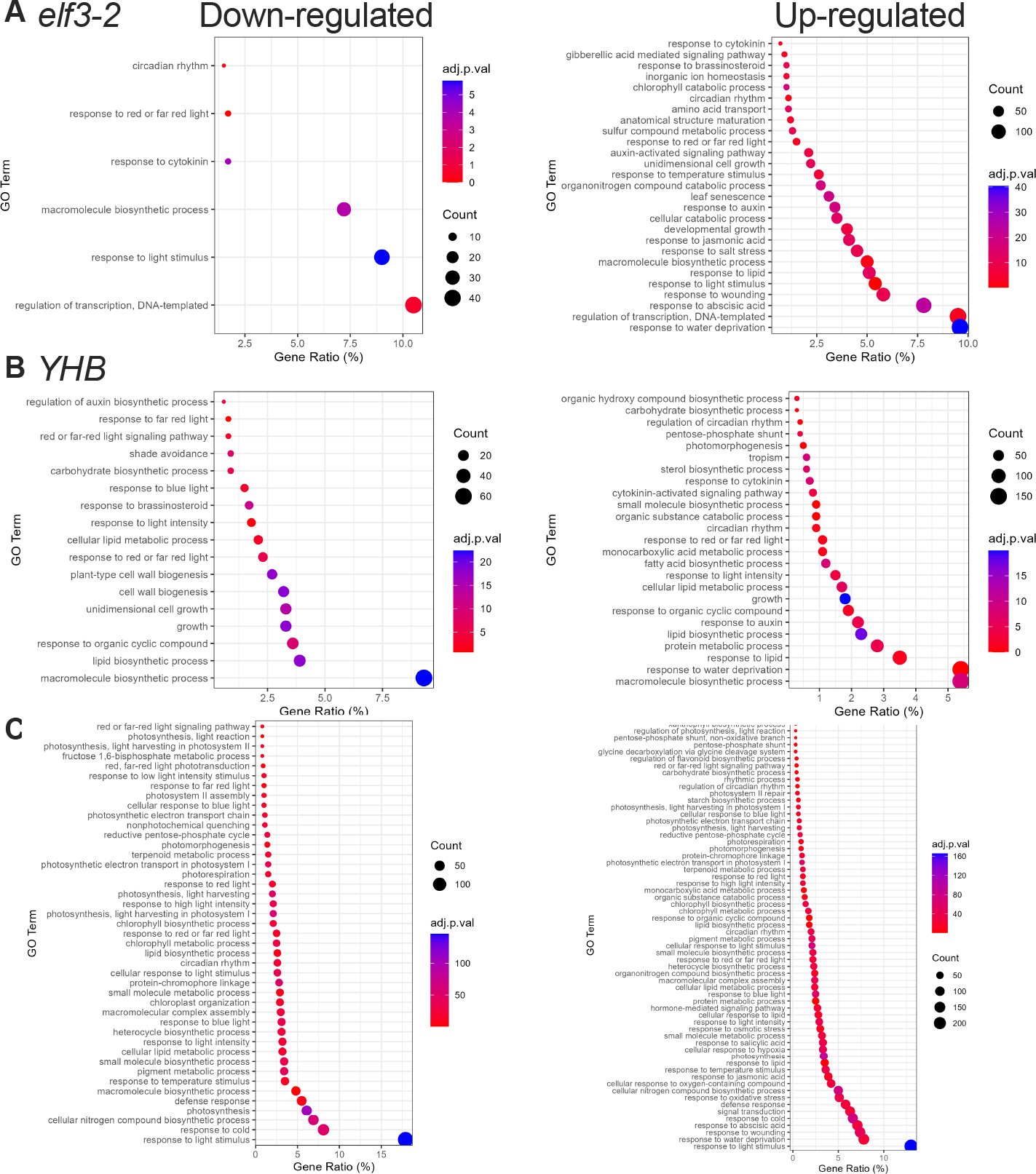
GO term enrichment in *YHBelf3* seedlings at ZT60 (a-c) Significantly enriched GO terms in *elf3-2* (A), *YHB* (B), and *YHBelf3-2* (C) seedlings compared to wild type at ZT60. Bar charts show number of transcripts mis-expressed in each GO term. Heat maps highlight significantly over- represented GO terms. See also Table S2.

**Figure S6.**
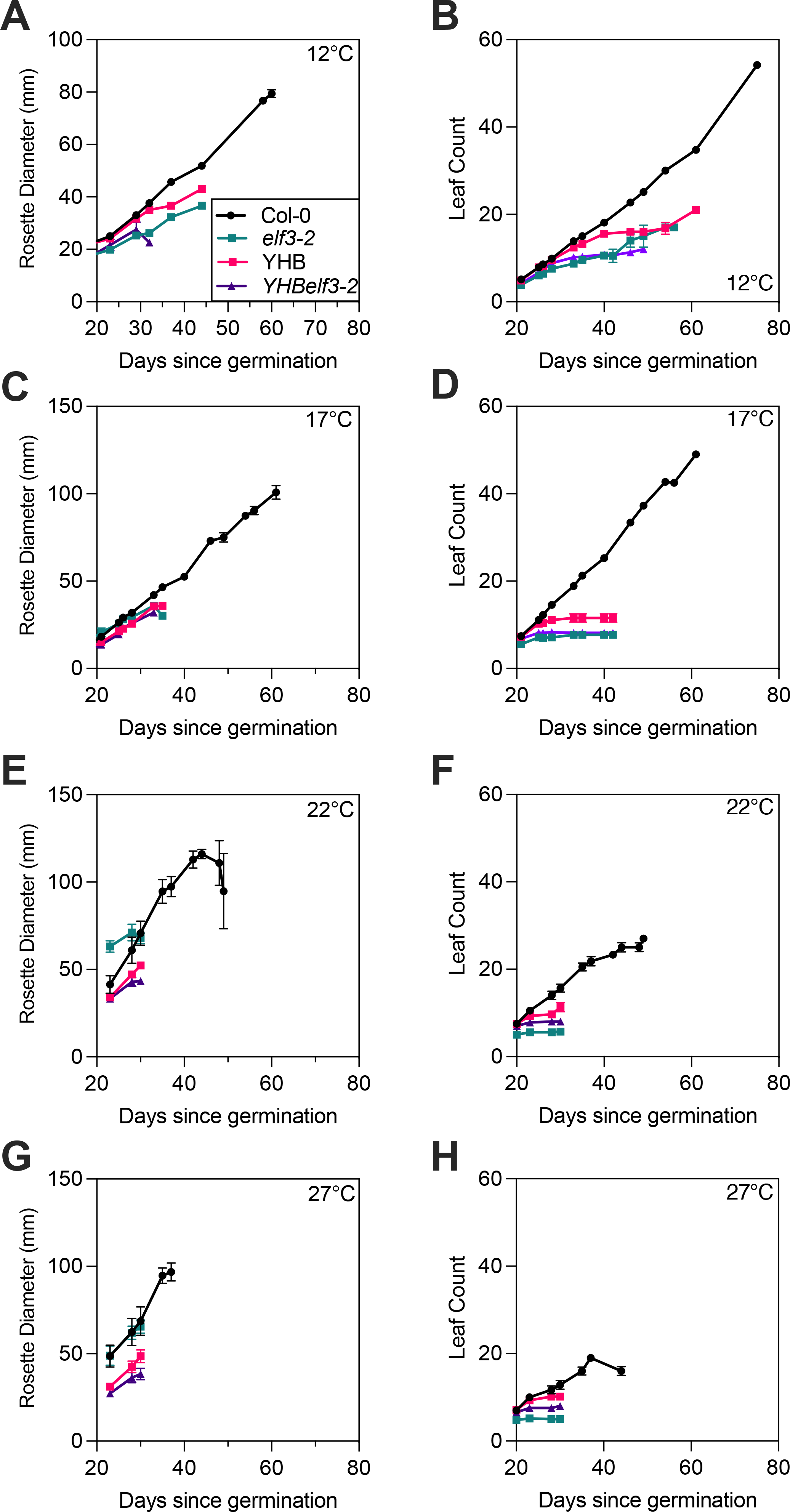
Ambient temperature has reduced effects upon *YHBelf3* growth throughout development. (A-D) Rosette diameter in wild type, *elf3-2*, *YHB*, and *YHBelf3-2* seedlings grown in 12:12 light:dark cycles at either 12°C (A), 17°C (B), 22°C (C), or 27°C (D). **(E-H)** Number of leaves in wild type, *elf3-2*, *YHB*, and *YHBelf3-2* seedlings grown in 12:12 light:dark cycles at either 12°C (E), 17°C (F), 22°C (G), or 27°C (H). Error bars indicate SEM.

Table S1. Differential gene expression in elf3, YHB, and YHBelf3 seedlings at ZT60. Summary statistics of transcripts mis-expressed compared to wildtype after 48 hours in constant darkness. Individual TPM for WT and each mutant for the given transcript is shown alongside Log2 fold change and adjusted p values. Transcript lists are separated into sheets for up and down-regulated transcripts for each mutant.

Table S2. GO term enrichment in elf3, YHB, and YHBelf3 seedlings at ZT60. List of GO terms for direct biological processes from DAVID for each list of significantly mis-expressed genes from Table S1 showing count of mis-expressed genes associated with the GO term, percentage of genes in the list associated with the GO term, p-value and Benjamini adjusted p-value for each GO term.

Table S3. Genomic loci associated with ELF3 and/or phyB. Data were re-analysed from (Jung et al., 2016, Ezer et al., 2017).

Table S4. Oligonucleotides used in this study.

