## Supplementary material for "Manipulation of Photosensory and Circadian Signalling Restricts Developmental Plasticity in Arabidopsis": Table S4

**Table S4. Oligonucleotides used in this study**

| **Gene** | **AGI** | **Orientation** | **Sequence** | **Reference:** |
| --- | --- | --- | --- | --- |
| *CCA1* | TAIR: AT2G46830 | Forward | CAGCTCCAATATAACCGATCCAT | ^1^ |
|  |  | Reverse | CAATTCGACCCTCGTCAGACA | ^1^ |
| *LHY* | TAIR: AT1G01060 | Forward | CAATGCAACTACTGATTCGTGGAA | ^1^ |
|  |  | Reverse | GCTATACGACCCTCTTCGGAGAC | ^1^ |
| *GIGANTEA* | TAIR: AT1G22770 | Forward | ACTAGCAGTGGTCGACGGTTTATC | ^2^ |
|  |  | Reverse | GCTGGTAGACGACACTTCAATAGATT | ^2^ |
| *PHYB* | TAIR: AT2G18790 | Forward | GACGTCGGAGGAGGATAGAA | This study |
|  |  | Reverse | AAACAAAAGTCTTGAAATTCTCTACAA | This study |
| *PHYB::YHB* |  | Forward | GACGTCGGAGGAGGATAGAA | This study |
|  |  | Reverse | GCCGCACTAGTGATTGAGC | This study |
| *35S::YHB* |  | Forward | CCCTGTACCTCGAAAGCGACCA | This study |
|  |  | Reverse | ACCCCGGTGAACAGCTCCTC | This study |
| *PRR9* | TAIR: AT2G46790 | Forward | GAAACAACGTTGGAGTAGAAGC | This study |
|  |  | Reverse | CTCTGGTACCGAACCTTTTTG | This study |
| *APX3* | TAIR: AT4G35000 | Forward | GCCGTGAGCTCCGTTCTCT | ^3^ |
|  |  | Reverse | TCGTGCCATGCCAATCG | ^3^ |
| *IPP2* | TAIR: AT3G02780 | Forward | GTATGAGTTGCTTCTGGAGCAAAG | ^3^ |
|  |  | Reverse | GAGGATGGCTGCAACAAGTGT | ^3^ |
| *APA1* | TAIR: AT1G11910 | Forward | CTCCAGAAGAGTATGTTCTGAAAG | ^3^ |
|  |  | Reverse | TCCCAAGATCCAGAGAGGTC | ^3^ |
| *CER* | *pDAN0869* | Forward | CTTGTACAAAGTGGTGCGAATGG  TGAGCAAGGGCGAGG | This study |
|  | *pDAN0870* | Reverse | GGCTCCAGCTTCCACCCCTAGACTTGTA  CAGCTCGTCCA | This study |

1.	Mockler, T. C. Regulation of flowering time in Arabidopsis by K homology domain proteins. *Proc Natl Acad Sci* **101**, 12759-12764 (2004).

2.	Mizuno, T. et al. The EC night-time repressor plays a crucial role in modulating circadian clock transcriptional circuitry by conservatively double-checking both warm-night and night-time-light signals in a synergistic manner in Arabidopsis thaliana. *Plant & Cell Physiology* **55**, 2139-2151 (2014).

3.	Nusinow, D. A. et al. The ELF4-ELF3-LUX complex links the circadian clock to diurnal control of hypocotyl growth. *Nature* **475**, 398-402 (2011).
